# Mechanistic evidence of widespread insecticide resistance among Illinois West Nile virus vectors (*Culex pipiens* and *Culex restuans*)

**DOI:** 10.1101/2024.09.17.613396

**Authors:** Kylee R. Noel, Chang-Hyun Kim, Chris M. Stone

## Abstract

**Background:** Mosquitoes are major vectors of arboviruses and other vector-borne diseases, making them a significant public health concern worldwide. Mitigation of arboviral outbreaks relies largely on the use of insecticides, but the effectiveness of such responses is threatened by the evolution of insecticide resistance. Monitoring mosquito susceptibility to different insecticides is therefore vital for informed decisions regarding outbreak responses. In this study, we elucidate the patterns of resistance to two insecticide classes within the primary vectors of West Nile virus in the northeast and midwestern regions of the continental United States, *Culex pipiens* and *Culex restuans*.

**Methodology/Principal Findings:** Egg collections were performed throughout Illinois from 2018-2020, and adults were tested for insecticide resistance to permethrin and malathion. Individuals from each sampling location were sequenced to determine the presence of *kdr* target-site mutations, and biochemical assays were performed to determine increases in detoxification enzymes and insensitive acetylcholinesterase. Results from the bottle assays indicate variable resistance rates in Illinois, however lowered mortality was found in most of the regions that were tested. The *kdr* mutation (L1014F) was present in 50% of *Culex pipiens* sequenced, and more prevalent in southern Illinois compared with northern and central (*p* < 0.001). Different mechanisms were predictive of resistance by species and insecticide, with permethrin resistance being affected by *kdr*-allele frequency and oxidase levels and malathion resistance by *α*- and *β*-esterases in *Cx. pipiens*. For *Cx. restuans α*-esterase and oxidase levels were predictive of permethrin resistance while *β*-esterase and insensitive acetylcholinesterase levels were predictive of malathion resistance.

**Conclusions/Significance:** We documented variation in insecticide resistance levels that appear to be driven by population differences in *kdr* mutation rates and metabolic resistance mechanisms. The presence of different mechanisms in species and regions has implications for approaches to resistance management and highlights the need to implement and maintain insecticide resistance monitoring practices.

**Author Summary:** Mosquitoes are the vectors of many major diseases including malaria, dengue, yellow fever, zika, and West Nile virus. Insecticides are often used to control mosquitoes and the outbreaks they cause. However, evidence has shown that populations of different mosquito species worldwide have developed resistance to our most common insecticides. This study shows that West Nile virus vectors in Illinois, (*Culex pipiens* and *Culex restuans*) are no exception to this trend. Egg collections were made throughout the state during the 2018-2020 field seasons and the resulting adults were tested for resistance to two common insecticides using the CDC’s bottle bioassay protocol. The results indicate that rates of resistance vary throughout the state and population differences in resistance mechanisms are driving this variation.

## Introduction

Vector-borne diseases are a global public health threat that account for more than 17% of all infectious diseases and caused an estimated 847,472 deaths in 2021 alone [1,2]. West Nile virus (WNV) is an important disease, primarily transmitted by *Culex* mosquitoes, that causes symptoms ranging from mild fever to severe, lethal neuroinvasive disease and is endemic in Africa, Europe, the Middle East, North America, and West Asia [3]. WNV is the leading cause of mosquito-borne disease in the continental United States, with a total of 51,702 cases reported to the Centers for Disease Control and Prevention (CDC) between 1999 and 2019; however, some studies have shown that this is likely an underestimation [4,5]. This underestimation is due to many factors, including asymptomatic cases or cases with mild symptoms that are not severe enough for the individual to seek medical care. People experiencing socioeconomic hardships who lack the resources to obtain medical attention, and who may be especially vulnerable to infection, also likely contribute to underreporting [4,6].

There are currently no vaccines for prevention or specific medications to treat WNV in people. However, transmission of WNV can be limited by controlling mosquito populations which is done by use of proactive mosquito control programs, typically employing larval control methods [3,7]. Reactive or emergency control methods are typically initiated when surveillance detects high levels of enzootic infection or the reporting of human cases [7]. Currently, the most effective reactive response is the use of adulticides, which are widely used to reduce vector populations by public health departments and mosquito control districts during an outbreak [8,9]. There is concern that the amount of insecticides used, not only in vector control but for instance, also for agriculture, could exert a strong selective pressure for resistance and reduce the effectiveness of mosquito adulticides [10,11].

Resistance to the most common classes of insecticides used in vector control, pyrethroids, and organophosphates, have been found in *Culex* populations around the world [12,13]. There are several different mechanisms found in these populations that limit the effects of the insecticides and mosquito populations can exhibit more than one mechanism at a time [14–17]. Two major resistance mechanisms are target-site mutations and metabolic resistance [18,19]. A target-site mutation occurring in the sodium channel gene results from a single nucleotide polymorphism (SNP) changing a Leucine to either a Phenylalanine (L1014F) or Serine (L1014S) and confers resistance to pyrethroids. This form of resistance is called knockdown resistance (*kdr*) because it allows the mosquito to avoid the knockdown effect of the insecticide [13]. Another important target-site mutation is caused by a SNP in the *ace*-1 gene encoding acetylcholinesterase (ACHE), which is the target of organophosphate insecticides [20]. This mutation is due to a replacement of Glycine by Serine (G119S), with the result being ACHE that is insensitive to the insecticide [21]. Metabolic resistance occurs when insects can sequester, metabolize, or detoxify insecticides more efficiently through amplification of detoxification genes or the overexpression of these genes. Two major detoxification gene families involved in this process are cytochrome P450s and carboxylesterases [22].

Insecticide-based strategies are the most readily implemented tools to control outbreaks of mosquito-borne disease on a global scale [23]. As insecticides are crucial for these emergencies, and the panel of available insecticides is limited, it is important to monitor and manage resistance in mosquito populations [24]. Phenotypic resistance to permethrin, a pyrethroid, and malathion, an organophosphate, have been documented in *Culex* populations in Illinois [25]. However, the degree of resistance throughout the rest of the state and the underlying resistance mechanisms of these *Culex* populations are unknown. We must understand the prevalence and distribution of resistance to have a plan of action when a mosquito-borne disease outbreak occurs. To this end, surveys of *Culex* mosquito populations throughout Illinois were taken throughout 3 sampling seasons to detect phenotypic resistance using CDC bottle bioassays, and molecular methods were employed to determine the genetic and enzymatic contributions.

## Methods

### Sample Collections

Locations were chosen throughout the state that were of public health interest in cooperation with Illinois local public health departments and mosquito abatement districts, usually located near West Nile virus sentinel traps. At each sampling location, 5-gallon bins filled with 2 gallons of grass infusion were set out overnight to attract gravid *Culex* mosquitoes to oviposit. Bins were checked the following day for the presence of egg rafts. A small paint brush was used to gently lift the rafts out of the water, and they were then placed into 12-well tissue culture plates filled with DI water for transportation back to the lab. Each egg raft was individually placed into a well and identifications were performed at the first larval instar to determine the species as either *Culex pipiens* or *Culex restuans* [26,27].

### Larval Rearing and Adult Maintenance

After identification, batches of 200 larvae of the same species were transferred to white enameled pans containing 1.5 L of DI water. The larvae were fed a diet of Tetramin (Tetra Holding (US)), brewer’s yeast (MP Biomedicals LLC), and rabbit chow (Kaytee Products, Inc.) in a 1:1:1 mixture. Pans received 75 ± 1 mg of diet on days 0-4, then 100 ± 1 mg daily starting on the 5th day. Water changes were performed as needed. Pupae were separated from larvae daily and placed in a container of fresh DI water within adult cages for emergence. Adults were provided with flasks containing 10% honey solution with dental roll wicks. Pans and adult cages were contained in environmental chambers under standard insectary conditions (26 ± 1⁰C, 70 ± 8% relative humidity, and 16 L:8 D photoperiod).

### Phenotypic resistance assays

The resistance status of female mosquitoes aged 3-5 days was tested using CDC bottle bioassays [28]. Each assay consisted of four 250 mL Wheaton bottles, coated with 1 mL of the respective insecticide mixed with acetone, and 1 control bottle coated with 1 mL of acetone only. About 25 females were introduced into each bottle and were then observed over 2 hours, tracking mortality. Each location was tested for permethrin and malathion resistance using this method, except in the 2019 sampling season when only permethrin was tested. We used established diagnostic times for *Cx. pipiens* of 30 minutes for permethrin and 45 minutes for malathion for both species. A colony of susceptible *Cx. pipiens* mosquitoes started from the CDC’s “Chicago” strain was also used to confirm these diagnostic times. The ratio of mosquitoes that have not been knocked down or killed after that time was used to determine resistance levels. An average across experimental bottles of less than 90% mortality at the diagnostic time is considered indicative of resistance to that insecticide [28]. Tested mosquitoes were frozen immediately after completion of the assay at -80°C for further processing.

### Detection of *kdr* point mutations

A subset of 10 mosquitoes from each collection location were selected for sequencing of the *para-*type voltage-gated sodium channel for the *kdr* point mutation (L1014F/S). Mosquitoes were placed into individual tubes with 5 stainless steel beads, 100 µl buffer BE, 40 µl buffer MG, and 10 µl liquid proteinase K (buffers from Takara Bio) and were homogenized using the TissueLyser II (Qiagen). DNA was extracted using the NucleoSpin 96 DNA RapidLyse Kit according to the manufacturer’s protocol (Takara Bio). After extraction, DNA concentrations were measured using Qubit dsDNA High Sensitivity assay on a Qubit 4 Fluorometer (ThermoFisher Scientific). Next, a PCR was run to amplify the 176 bp area of interest using the following primers from Chen et al.: forward 5ʹ - GTGTCCTGCATTCCGTTCTT -3’ and reverse 5ʹ - TTCGTTCCCACCTTTTCTTG-3’ [29]. The primers were modified to include the Illumina overhang adapter sequences. Each reaction contained 12.5 µl of Q5® High-Fidelity 2X Master Mix (New England BioLabs), 1.25 µl of each 10 µM forward and reverse primer (Integrated DNA Technologies), 8 µl of water, and 2 µl of target sample for a total reaction volume of 25 µl. The reaction comprised of 1 cycle at 95°C for 5 min., 40 cycles at 95°C for 30 sec., 52°C for 30 sec., and 72°C for 1 min. with a final extension step of 72°C for 5 min. Next, samples with multiple banding were run through gel cassettes using the PippinHT (Sage Science) to size select for the band of interest.

PCR purification was performed on the DNA band selected from the PippinHT using the Mag-Bind® Total Pure NGS kit according to the manufacturer’s protocol (Omega Bio-Tek). A 1.8x ratio (45 µl) of Mag-Bind® Total Pure was used with 25 µl of PCR product. Next, an index PCR was performed on the purified product to attach dual indices and Illumina sequencing adapters using the Nextera XT Index Kit. Each reaction contained 25 µl of Q5® High-Fidelity 2X Master Mix (New England BioLabs), 4 µl of Nextera UD Indexes for labeling, 16 µl of water, and 5 µl of target sample for a total reaction volume of 50 µl. The reaction comprised of 1 cycle at 95°C for 3 min., 12 cycles at 95°C for 30 sec., 55°C for 30 sec., and 72°C for 30 sec. with a final extension step of 72°C for 5 min. Another PCR purification was performed on the indexed samples using a 0.85x ratio (42.5 µl) of Mag-Bind® Total Pure with 50 µl of product. Libraries were quantified on a Qubit 4 Fluorometer and were then pooled in equal amounts according to product concentration for sequencing. Samples were sequenced using the Illumina MiSeq Nano V2 platform at the W. M. Keck Center for Comparative and Functional Genomics at the University of Illinois at Urbana-Champaign. The libraries were sequenced from both ends of the molecules to a total read length of 176 nt.

Preprocessing, mapping, and SNP calling of the next-generation sequencing (NGS) data was performed using Geneious Prime v2023.1.2. Data were imported, paired, and trimmed using the BBDuk trimmer plugin. Next, overlapping paired reads were merged into single reads using BBMerge, duplicate reads were removed using Dedupe, and chimeric reads were filtered from sequencing data using UCHIME. The reads were mapped to a reference sequence (GenBank: AY283036.1) and SNPs were identified using the SNPs per sample Geneious workflow.

### Biochemical Assays

A subset of mosquitoes from each location were randomly selected to test for increased enzyme activity using microplate enzyme assays. Five assays were performed for each mosquito: protein, elevated non-specific *α*-esterase, elevated non-specific *β*-esterase, mixed function oxidase (MFO), and an insensitive acetylcholinesterase (ACHE) assay. Excluding the MFO, these assays directly quantify the enzyme activity within individual mosquitoes. Cytochrome P450s are primarily associated with heme in non-blood-fed mosquitoes, so the MFO assay indirectly estimates cytochrome P450 activity from the heme content within each mosquito [30]. The assay procedures were followed as in McAllister et al. [31], each sample was run in triplicate per assay and absorbance values were read using a BioTek 800 TS absorbance reader (Agilent, Santa Clara, CA). Adult female mosquitoes were kept on ice and homogenized individually in 100 µl of potassium phosphate (KPO_4_) buffer. The mosquito homogenate was diluted to 2 ml with additional KPO_4_ buffer to yield enough material to run all assays. All chemicals were purchased from Sigma-Aldrich Chemical Co. (St. Louis, MO) unless otherwise noted.

The protein assay measures the amount of total protein present and is used to correct for size when comparing different individuals. This assay was performed using 10 µl of mosquito homogenate and 200 µl of Bradford reagent solution (ThermoFisher Scientific) per well. The samples were mixed thoroughly, incubated at room temperature for 5 minutes, and the absorbance was measured at 595 nm. For the *α* and *β* non-specific esterases 100 µl of either *α*-naphthyl acetate or *β*-naphthyl acetate was mixed with 100 µl of mosquito homogenate and incubated at room temperature for 40 minutes. After incubation, 100 µl of Dianisidine solution was added to each well and incubated for an additional 8 minutes at room temperature, then the absorbance was measured at 540 nm. The MFO assay was performed using 100 µl of mosquito homogenate, 200 µl of the substrate 3,3’,5,5’-tetramethybenzidine, and 25 µl of 3% hydrogen peroxide in each well. The plates were incubated at room temperature for 20 minutes and the absorbance was read using the 620 nm filter. The ACHE assay combined propoxur with the substrate acetylthiocholine iodide (ATCH) (100 µl), mixed with 100 µl of mosquito homogenate. Then 100 µl of 5,5’ Dithio-bis-2-nitrobenzoic acid was added to each well. Absorbances were read immediately (T = 0 min.) at 414 nm, then the plates were covered and refrigerated overnight, and a second reading was taken at T = 24 hours. Absorbance was read at different time points because it is a kinetic assay. The value used for statistical analysis was obtained by subtracting the T = 0 min. reading from the T = 24 hours reading.

### Data Analysis

All data analyses were performed in R version 4.3.2 [32]. Maps indicating mortality at the diagnostic time were produced using data from the CDC bottle bioassays using the package ‘*ggmap*’ [33]. Differences in survival from the bottle assays, compared with a susceptible control strain of *Cx. pipiens* were analyzed with a Cox proportional-hazards model using a robust variance with the package ‘*survival’* [34]. Hazard ratios were calculated to determine the probability of mortality (i.e., hazard) compared to the control strain. Species differences in mortality to each insecticide were assessed using the ‘*ANOVA’* procedure within the package ‘*car’*, using a type III hypothesis [35]. A map displaying the genotypes of the *kdr* SNP by county was created using the NGS results. A comparison of F allele frequency in Northern, Central, and Southern populations was performed using the ‘*ANOVA’* procedure followed by a *post hoc* pairwise test of means using the Tukey method (package ‘emmeans’) [36]. Results from the biochemical assays were analyzed by region using a Kruskal-Wallis test (‘kruskal.test’ from the ‘stats’ package) [32]. To determine significant differences from the control strain, Dunn’s (1964) test of multiple comparisons was performed using a Bonferroni correction. We analyzed the susceptibility to permethrin and malathion of *Cx. pipiens* and *Cx. restuans* using mortality at the diagnostic time as the response variable for each insecticide and examined the effects of five explanatory variables: F allele frequency, *α*- and *β*-esterase concentration, MFO concentration, and ACHE insensitivity. The best-fitting linear model for each species with each insecticide was determined by the Akaike Information Criterion (AIC) using the function ‘*aictab*’ within the package ‘*AICcmodavg*’ [37].

## Results

### Phenotypic resistance

In total 44 sites were sampled during the 2018-2020 collection seasons. *Cx. pipiens* were collected from 36 of the sites and *Cx. restuans* were collected from 8 of the sites. Results from CDC bottle bioassays testing for permethrin and malathion were highly variable throughout the state (Fig. 1). Average mortality at the diagnostic time was also calculated for each insecticide by region (Table 1 & 2). *Cx. pipiens* from all regions had an average mortality of less than 90% at the 30-minute diagnostic time for permethrin indicating resistance. Of the four regions in which *Cx. restuans* populations were tested using permethrin, only the West Chicago region displayed resistance. For malathion, only the Marion region *Cx. pipiens* displayed resistance, and all other populations tested had an average mortality greater than 90% at the 45-minute diagnostic time.

**Fig. 1.**
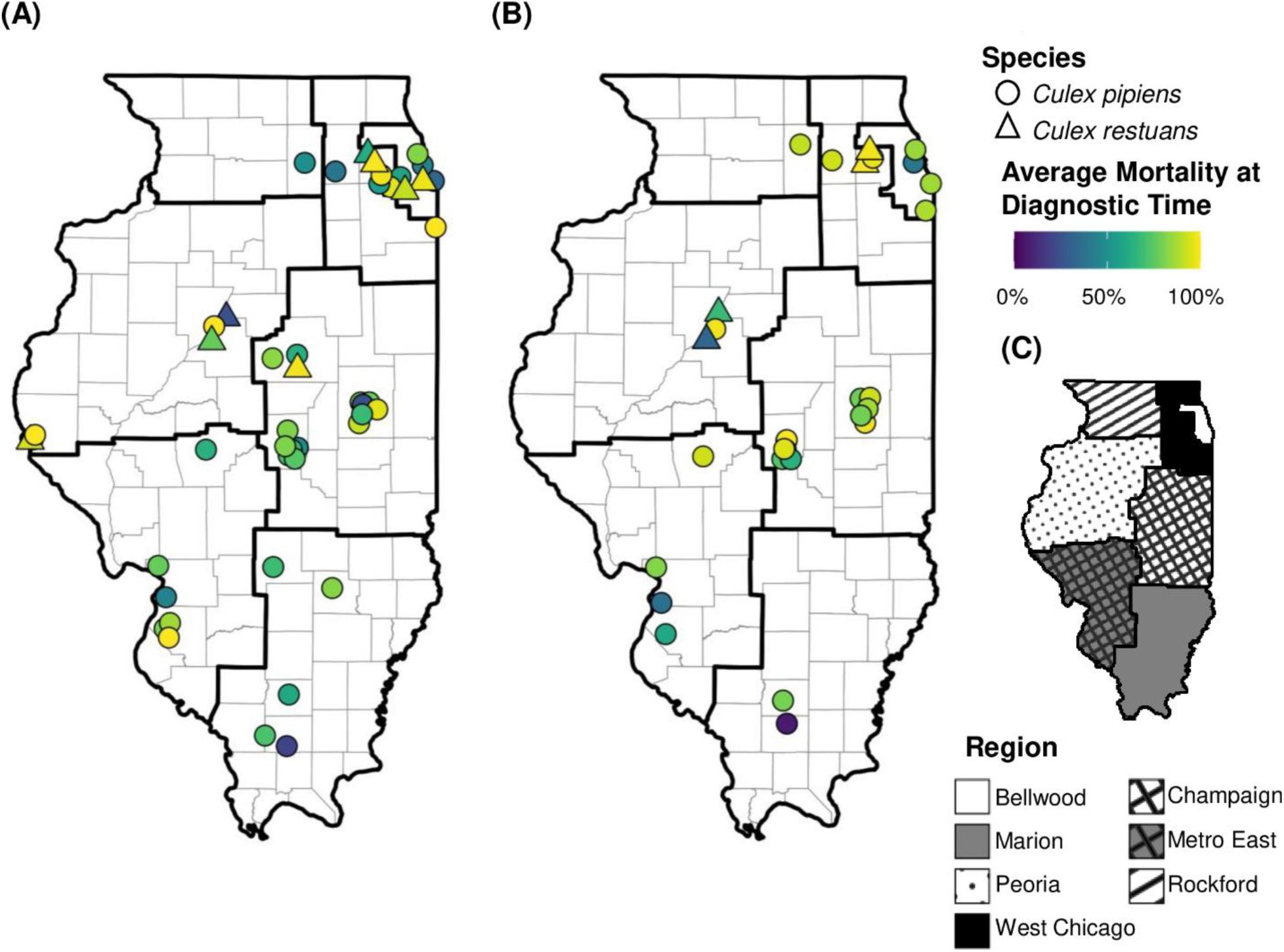
Susceptibility to permethrin and malathion is highly variable throughout IL. (A) Results from CDC bottle bioassay testing permethrin with mortality reported at the diagnostic time of 30 minutes. (B) CDC bottle bioassay results for malathion with mortality reported at the diagnostic time of 45 minutes. (C) Map depicting the division of Illinois into the 7 regions used in this study.

**Table 1.** Survival analysis by region for permethrin.

| Region | *Culex pipiens* |  |  |  | *Culex restuans* |  |  |  |
| --- | --- | --- | --- | --- | --- | --- | --- | --- |
|  | Mortality<sup>1</sup> | HR<sup>2</sup> | 95% CI | p-value | Mortality<sup>1</sup> | HR<sup>2</sup> | 95% CI | p-value |
| Bellwood | 64% | 0.08 | 0.05, 0.12 | <0.001 | 95% | 0.18 | 0.11, 0.29 | <0.001 |
| Champaign | 73% | 0.10 | 0.06, 0.14 | <0.001 | 98% | 0.23 | 0.13, 0.42 | <0.001 |
| Marion | 63% | 0.06 | 0.04, 0.09 | <0.001 | - | - | - | - |
| Metro East | 74% | 0.09 | 0.006, 0.13 | <0.001 | - | - | - | - |
| Peoria | 63% | 0.59 | 0.38, 0.92 | 0.021 | 99% | 0.04 | 0.02, 0.07 | <0.001 |
| Rockford | 50% | 0.08 | 0.05, 0.11 | <0.001 | - | - | - | - |
| West Chicago | 69% | 0.11 | 0.07, 0.16 | <0.001 | 72% | 0.05 | 0.03, 0.09 | <0.001 |

<sup>1</sup>Average mortality at the diagnostic time (30 minutes), <sup>2</sup>Hazard Ratio; \* $p$ values were calculated using the Cox-proportional hazard model.

Hazard ratios comparing mosquito survival to permethrin in each region were significantly different from the control strain for all regions and both species (Table 1). For malathion, *Cx. pipiens* from all regions except Peoria (*p* = 0.3), were significantly different from the control strain (Table 2). Only two regions were tested for *Cx. restuans* malathion susceptibility and both were significantly different from the control strain. However, the West Chicago population had a hazard ratio greater than one meaning that this population was more likely to die by the diagnostic time compared to the control strain. *Cx. pipiens* had greater variation in average mortality at the diagnostic time for both insecticides compared to *Cx. restuans*, and there was a significant difference in permethrin mortality at the diagnostic time between species (*p* = 0.0098) (Fig. 2).

**Fig. 2.**
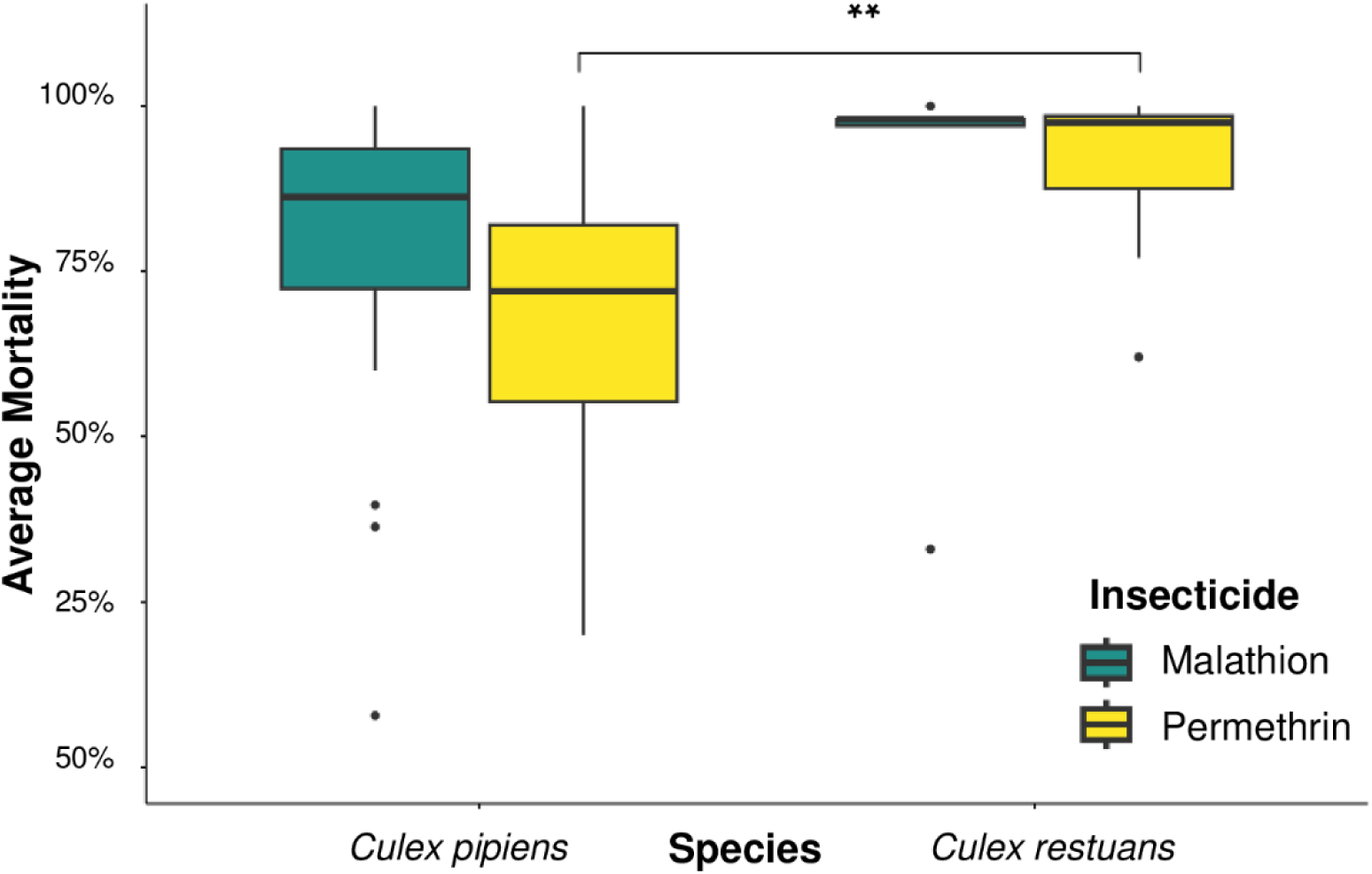
Difference in species average mortality at the diagnostic time from CDC bottle bioassays testing permethrin (yellow) and malathion (teal) susceptibility. There is a significant difference between *Cx. pipiens* and *Cx. restuans* average permethrin mortality. (** *p* < 0.01).

**Table 2.** Survival analysis by region for malathion.

| Region | *Culex pipiens* |  |  |  | *Culex restuans* |  |  |  |
| --- | --- | --- | --- | --- | --- | --- | --- | --- |
|  | Mortality<sup>1</sup> | HR<sup>2</sup> | 95% CI | p-value | Mortality<sup>1</sup> | HR<sup>2</sup> | 95% CI | p-value |
| Bellwood | 95% | 0.21 | 0.16, 0.28 | <0.001 | - | - | - | - |
| Champaign | 99% | 0.35 | 0.27, 0.45 | <0.001 | - | - | - | - |
| Marion | 79% | 0.11 | 0.08, 0.14 | <0.001 | - | - | - | - |
| Metro East | 95% | 0.19 | 0.15, 0.25 | <0.001 | - | - | - | - |
| Peoria | 100% | 0.81 | 0.56, 1.17 | 0.3 | 96% | 0.08 | 0.06, 0.12 | <0.001 |
| Rockford | 100% | 0.44 | 0.32, 0.62 | <0.001 | - | - | - | - |
| West Chicago | 100% | 0.51 | 0.39, 0.67 | <0.001 | 100% | 1.39 | 1.02, 1.88 | 0.036 |

<sup>1</sup>Average mortality at the diagnostic time (45 minutes), <sup>2</sup>Hazard Ratio; \* $p$ values were calculated using the Cox-proportional hazard model.

### Next-Generation Sequencing

Successful sequencing was performed on 511 total individuals, with 411 being *Cx. pipiens* and 100 *Cx. restuans.* None of the sequenced *Cx. restuans* showed a *kdr* point mutation. The L1014F mutation was present in a portion of the *Cx. pipiens* that were sampled, but not the L1014S mutation (Fig. 3A). Half of the sequenced *Cx. pipiens* had the F allele present, with 12% being homozygous for the mutation (Fig. 3B). The frequency of the F allele was higher in populations sampled in the southern part of the state compared with the northern (*p* < 0.0001) and central (*p* < 0.0001) portions of the state and was also significantly different between northern and central IL. (*p* < 0.05) (Fig. 3C).

**Fig. 3.**
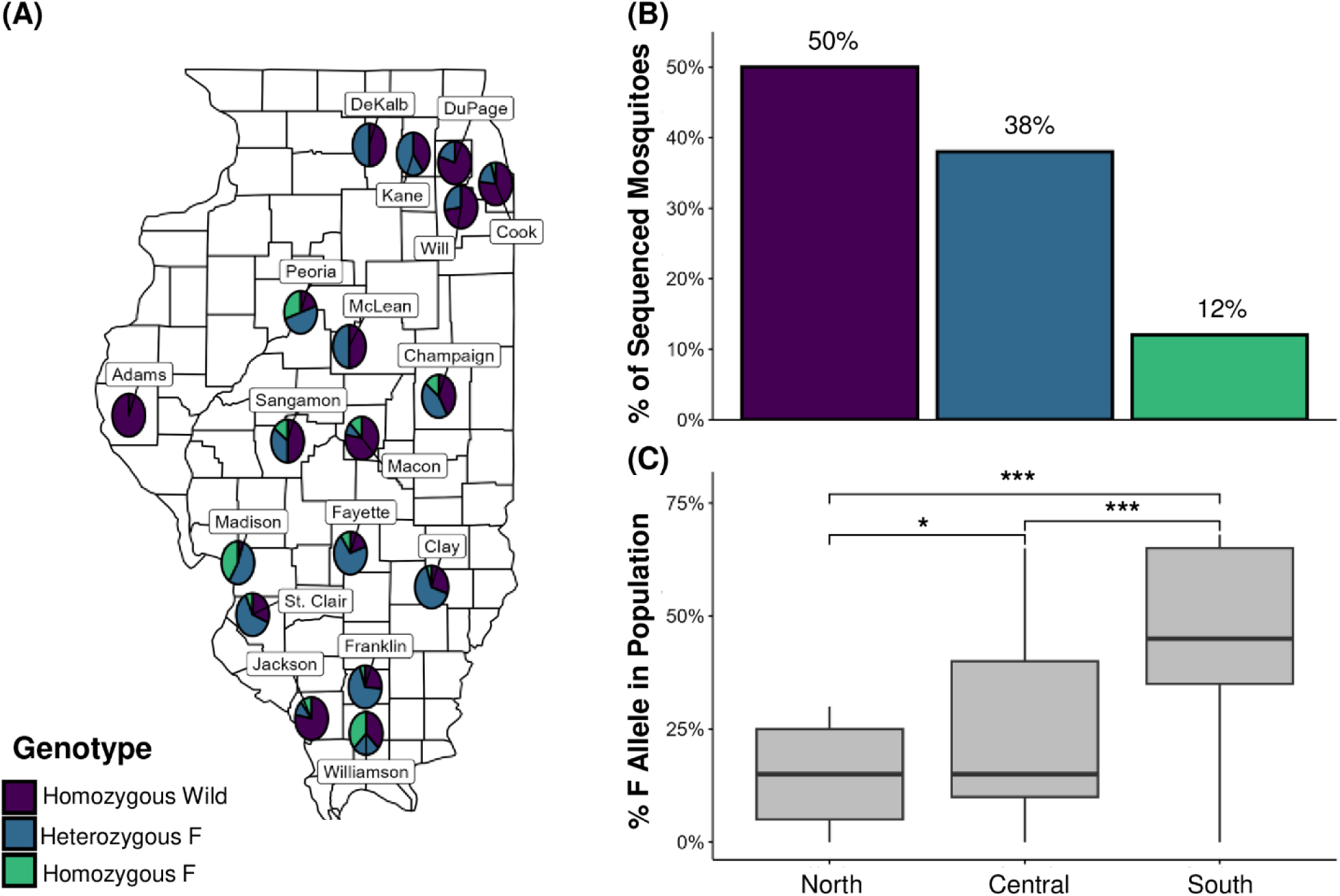
The *kdr* point mutation was detected throughout IL. in *Culex pipiens* and occurred at a higher frequency in southern IL. (A) Genotype frequencies for the *kdr* SNP (L1014F) of *Culex pipiens* sampled by county in IL. (B) Genotype frequency totals for all sequenced *Culex pipiens*. (C) Frequency of the F allele in the north, central, and southern parts of the state. The F allele frequency in the south is significantly different from the north and central parts of IL. (\*\*\**p* < 0.001); and the F allele frequency in the north is significantly different from central IL. (\**p* < 0.05).

### Biochemical Assays

A total of 383 individuals were used in the biochemical assays. Average results by region for each species can be found in Table 3 and Table 4. *α*-esterase levels were elevated in *Cx. pipiens* populations in the Champaign, Marion, Metro East, and Peoria regions, and in *Cx. restuans* populations from Bellwood and Champaign. *β*-esterase levels were not significantly different for any *Cx. pipiens* population compared to the control, however *Cx. restuans* populations from Peoria and West Chicago were lower than the control levels. MFO concentrations were elevated in *Cx. pipiens* populations from the Champaign, Marion, Peoria, and West Chicago regions, and elevated oxidase levels were found in *Cx. restuans* sampled from the Champaign and West Chicago regions. There were significant differences between the tested populations and the control in average absorbance from the ACHE assay.

**Table 3.** *Culex pipiens* biochemical assay results by region.

| Region | Average $\mu\text{g}$ α-esterase /mg Protein | Average $\mu\text{g}$ β-esterase /mg Protein | Average $\mu\text{g}$ oxidase /mg Protein | Average ATCH absorbance (nm) |
| --- | --- | --- | --- | --- |
| Bellwood | 55.93 | 22.35 | 4.57 | 0.08* |
| Champaign | 87.5* | 34.21 | 8.39* | 0.1* |
| Marion | 84.3* | 29.20 | 6.22* | 0.16 |
| Metro East | 74.95* | 25.72 | 5.91 | 0.12* |
| Peoria | 85.93* | 29.55 | 7.87* | 0.16 |
| Rockford | 51.55 | 22.43 | 5.61 | 0.14 |
| West Chicago | 61.80 | 23.85 | 6.54* | 0.10* |
| Control | 51.33 | 27.11 | 4.87 | 0.16 |

\*Significant difference from the control strain ( $p < 0.05$ ).

**Table 4.** *Culex restuans* biochemical assay results by region.

| Region | Average $\mu\text{g}$ α-esterase /mg Protein | Average $\mu\text{g}$ β-esterase /mg Protein | Average $\mu\text{g}$ oxidase /mg Protein | Average ATCH absorbance (nm) |
| --- | --- | --- | --- | --- |
| Bellwood | 75.94* | 28.52 | 5.51 | 0.07* |
| Champaign | 90.53* | 31.77 | 7.06* | 0.16 |
| Peoria | 60.26 | 22.51* | 4.78 | 0.08* |
| West Chicago | 57.49 | 22.28* | 6.75* | 0.09* |
| Control | 51.33 | 27.11 | 4.87 | 0.16 |

\*Significant difference from the control strain ( $p < 0.05$ ).

### Analysis of variation in resistance levels

Models were selected that best described the mortality at the diagnostic time for each combination of insecticide and species (Table 5). Each maximal model considered the five explanatory variables (F allele frequency, *α*- and *β*-esterase concentration, MFO concentration, and ACHE insensitivity), and the models were selected using the AIC. We found that *Cx. pipiens* permethrin mortality was best described by an interactive linear model using F allele frequency and oxidase as the explanatory variables. The F allele frequency in a population has a significant negative effect on average permethrin mortality, with mortality decreasing as the F allele frequency increases (Fig. 4A). Survival analysis showed that populations with medium or high frequency of the F allele differ significantly from populations with low F allele frequency, with low-frequency populations having an overall lower survival rate (Fig. 4B). The effect of oxidase alone is unclear and does not have a definite positive or negative effect on mortality. However, there is an interactive effect between F allele frequency and oxidase, where the effect of oxidase changes depending upon the frequency of the F allele. Malathion mortality in *Cx. pipiens* was best described by an additive linear model with *α*- and *β*-esterase as the explanatory variables. There was a negative effect of *α*-esterase on the average malathion mortality in *Cx. pipiens* populations, with increases in *α*-esterase concentration leading to lower average mortality or higher levels of resistance (Fig. 5A). There was also a negative effect of *β*-esterase concentration on the average malathion mortality in *Cx. pipiens* populations (Fig. 5B).

**Fig. 4.**
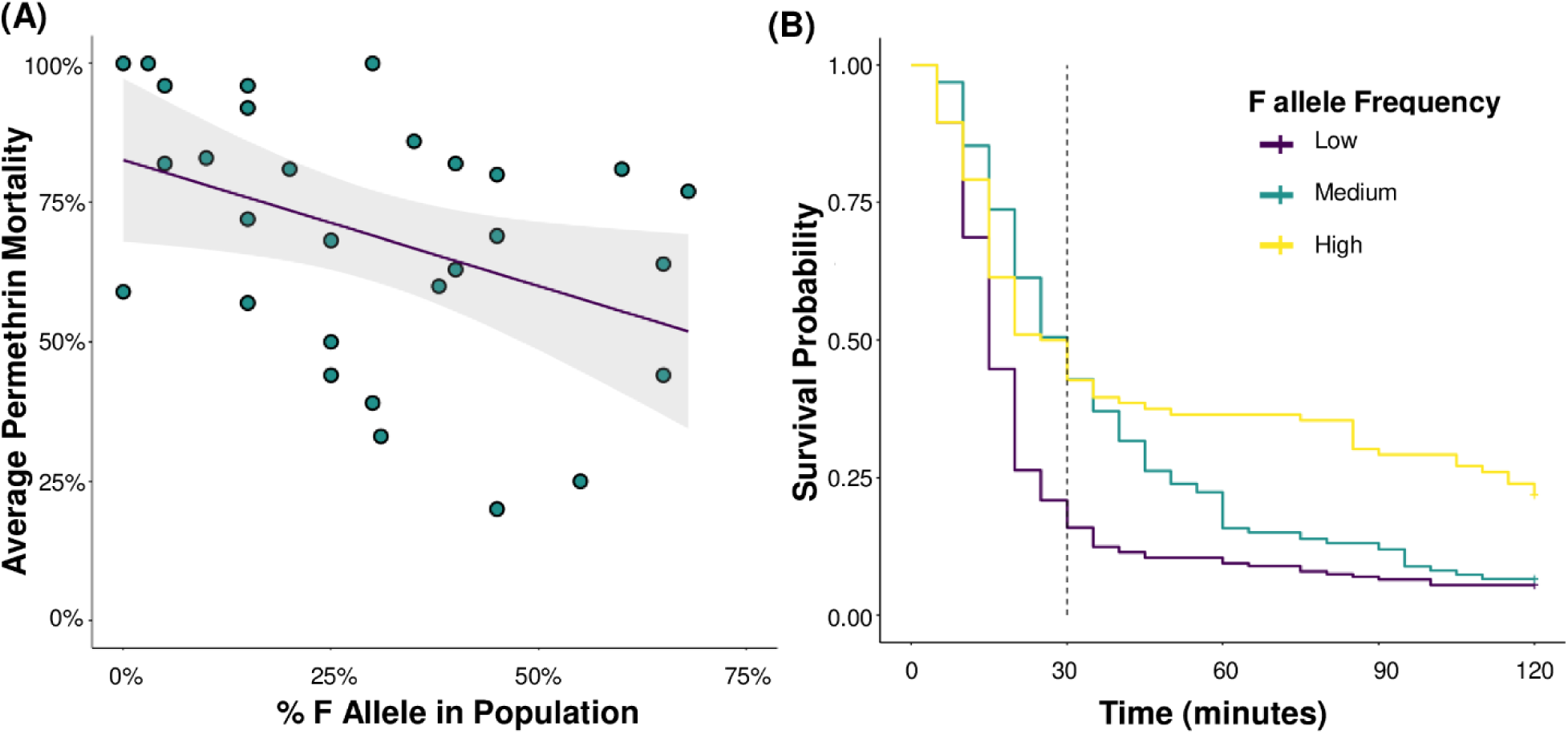
The effect of the *kdr* point mutation on *Culex pipiens* average permethrin mortality at the diagnostic time and survival probability. (A) There is a significant effect of F allele frequency on average permethrin mortality, with mortality decreasing (resistance increasing) with increasing F allele frequency (*** *p* < 0.001). (B) Survival analysis comparing populations with low, medium, and high F allele frequencies. Hazard ratios of high and medium frequency populations were significantly different from that of the low frequency populations (*** *p* < 0.001).

**Fig. 5.**
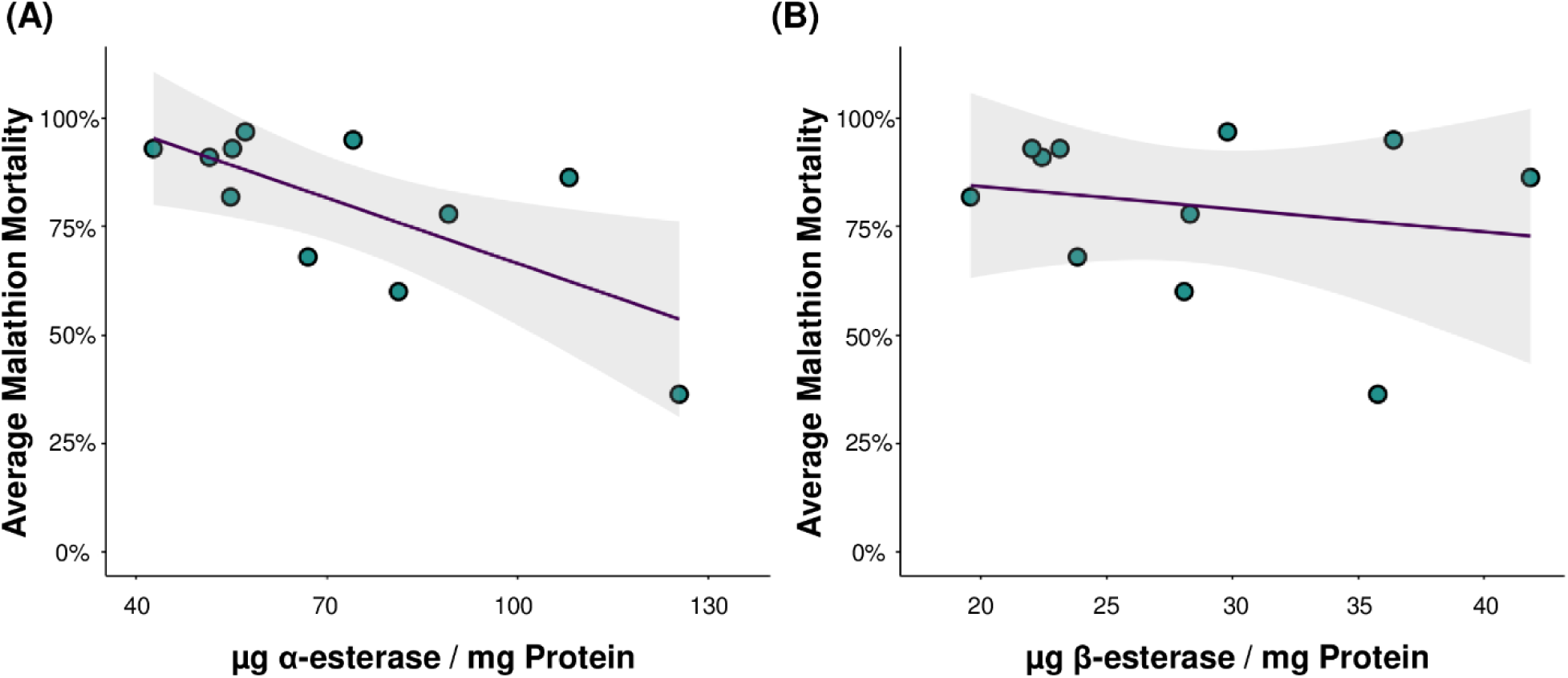
The effect of *α*- and *β*-esterase concentration on *Culex pipiens* average malathion mortality at the diagnostic time. (A) There is a significant effect of *α*-esterase concentration on malathion mortality, with mortality decreasing (resistance increasing), with increasing *α*-esterase concentration (*** *p <* 0.001). (B) There is also a significant effect of *β*-esterase concentration on malathion mortality, with mortality decreasing with increasing *β*-esterase concentration (*** *p <* 0.001).

**Table 5.**
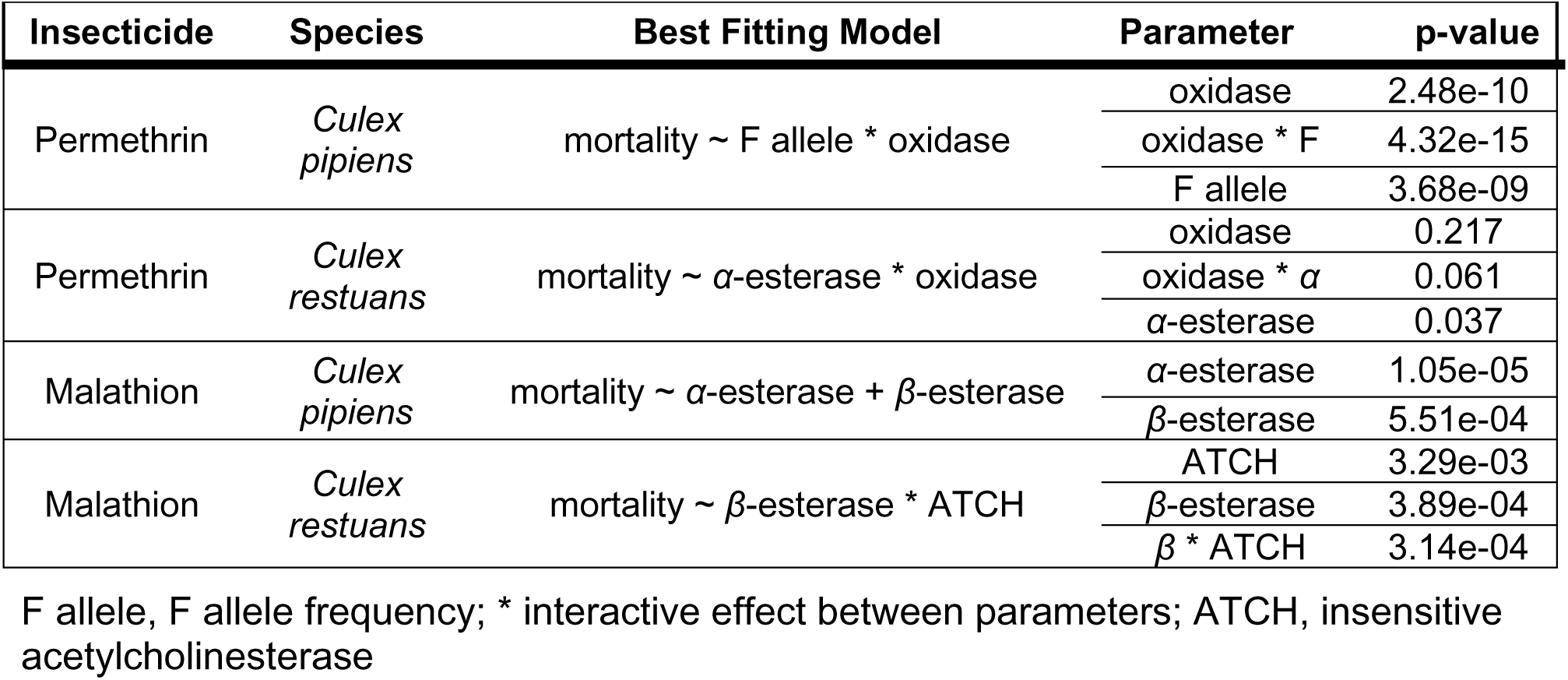
Best fitting linear models for insecticide resistance to permethrin and malathion for both species.

| Insecticide | Species | Best Fitting Model | Parameter | p-value |
| --- | --- | --- | --- | --- |
| Permethrin | *Culex pipiens* | mortality ~ F allele * oxidase | oxidase | 2.48e-10 |
|  |  |  | oxidase * F | 4.32e-15 |
|  |  |  | F allele | 3.68e-09 |
| Permethrin | *Culex restuans* | mortality ~ α-esterase * oxidase | oxidase | 0.217 |
| | | | oxidase * $\alpha$ | 0.061 |
| | | | α-esterase | 0.037 |
| Malathion | *Culex pipiens* | mortality ~ α-esterase + β-esterase | α-esterase | 1.05e-05 |
| | | | β-esterase | 5.51e-04 |
| Malathion | *Culex restuans* | mortality ~ β-esterase * ATCH | ATCH | 3.29e-03 |
| | | | β-esterase | 3.89e-04 |
| | | | $\beta$ * ATCH | 3.14e-04 |

F allele, F allele frequency; \* interactive effect between parameters; ATCH, insensitive acetylcholinesterase

Permethrin mortality in *Cx. restuans* populations was best described by an interactive linear model with α-esterase and oxidase as the explanatory variables. The best fit for *Cx. restuans* malathion mortality was an interactive linear model with *β*-esterase and insensitive ACHE as the explanatory variables of the model.

## Discussion

Understanding the presence and patterns of insecticide resistance in vector populations has important ramifications for public health. These patterns and the mechanisms associated with resistance can inform the choice of control methods and potentially help shed light on selective pressures and inform resistance management approaches. Thus, here we characterized insecticide resistance levels and investigated possible genetic and physiological mechanisms involved in resistance for *Cx. Pipiens* and *Cx. restuans* populations throughout the state of Illinois. Overall, we found that resistance to permethrin and malathion was highly variable throughout the state and that most regions have populations that are less susceptible to these insecticides when compared to a susceptible strain. We found evidence for multiple resistance mechanisms, including the presence of the L1014F mutation of the voltage-gated sodium channel and oxidase levels for permethrin and *α*- and *β*-esterase levels for malathion. Additionally, we found important differences between the two primary co-occurring West Nile virus vectors *Cx. pipiens* and *Cx. restuans*,

In the present study *Cx. pipiens* displayed greater variability in average mortality compared with *Cx. restuans*. A significant difference in the average mortality induced by permethrin at the diagnostic time was also found between these species, with *Cx. restuans* having higher mortality (greater susceptibility) compared with *Cx. pipiens.* As these species often co-occur in the same environments and share many ecological traits, these differences are somewhat surprising [38]. It is possible that there is a difference in selective pressures between the species. The application of insecticide to control mosquito populations is often in response to increases in reported human WNV cases which tend to occur during late July through early September [39]. This coincides with a temporal shift in *Culex* species, with *Cx. restuans* abundance decreasing in late July and *Cx. pipiens* becoming the predominant species through the rest of the season [40]. With this timing, *Cx. restuans* may not have the same level of exposure to insecticides as *Cx. pipiens.* This potential difference in selective pressure could be a contributing factor as to why the *kdr* mutation was only found in *Cx. pipiens* populations as well. Although fewer *Cx. restuans* were sequenced for the *kdr* mutation overall, previous work has also indicated the absence of the *kdr* point mutation in *Cx. restuans* populations in east-central Illinois [41]. The greater variation in resistance levels, as well as geographic variation in mechanisms that appear to play a role, that was observed in *Cx. pipiens* could possibly be due to *Cx. pipiens* belonging to a species complex with the variation coming from subtle differences between the subspecies [42]. Additional sampling work is needed to determine the exact distributions of these subspecies throughout IL. as there is evidence of hybridization zone expansion for *Cx. pipiens* and *Cx. quinquefasciatus,* from collected hybrids and updated estimates of geographical range maps [43,44]. We found that the *kdr* point mutation was more prevalent in the populations sampled in southern IL. compared with northern and central IL. Evidence shows that the kdr mutation rate varies between the species in the *Culex* complex [45]. However, additional work is needed to determine how this relates to IL. *Culex* populations. For instance, it is also possible that regional differences in pesticide use and resistance management practices in agriculture and public health could have selected for different resistance mechanisms, but more information on the use of insecticides in IL. is required to determine if this is a factor [46].

The model that best described *Cx. pipiens* permethrin mortality was an interactive linear model using F allele frequency and oxidase as the explanatory variables, which makes sense given that both variables are implicated in pyrethroid resistance [13,47,48]. There is a clear negative effect of *kdr* on mortality (Fig. 4), but the interaction with oxidase is not as straightforward. In populations with low frequencies of the F allele, individuals that also had increased levels of oxidases had higher mortality than individuals with lower oxidase levels. However, in populations with high frequencies of the F allele, individuals with increased oxidase levels had lowered mortality, which is what we would expect to see as *kdr* and P450 detoxification have been shown to exhibit a multiplicative interaction [47]. One explanation could be due to environmental differences between populations, as P450-mediated permethrin resistance has associated fitness costs that varies according to the environment [49]. Additionally it has been shown that there is plasticity in which P450s are selected for in different populations which could lead to varying fitness costs between populations [50]. A similar situation is seen when looking at the best fitting model for *Cx. restuans* permethrin mortality, where there is an effect of *α*-esterase leading to decreased mortality but looking at *α*-esterase and oxidase together does not give a clear picture. Additional testing of *Cx. restuans* populations would be necessary to clarify these relationships.

Looking at malathion mortality, the best fitting model for *Cx. pipiens* is an additive linear model with *α*- and *β*-esterase as the explanatory variables. Overproduction of nonspecific carboxylesterases is known to be involved in organophosphate, carbamate, and pyrethroid resistance in insects [51]. The most prevalent amplified esterase genes are *Est-3,* coding for *α*-esterase, and *Est-2*, which codes for *β*-esterase. Due to their close proximity these genes are often co-amplified and considered together as a single ‘super locus’ (*Ester)*, although *Est-2* tends to have a greater transcription [52]. Paton et al. found a 3:1 ratio of *β* to *α*-esterase from individual *Cx. quinquefasciatus* homogenates [52]. This differs from the levels that were found in this study, as *β*-esterase concentrations were not found to be significantly elevated compared to the control, whereas the *α*-esterase concentrations were elevated in some testing regions. *β*-esterase is also included in the best fitting model for *Cx. restuans* malathion mortality, along with insensitive ACHE. However, regional averages of ATCH absorbance tended to be significantly lower than the control strain. There are fitness costs associated with the resistant alleles on the *ace-1* and *Ester* loci which could explain the low concentrations of *β*-esterase and insensitive ACHE detected in the sampled *Cx. restuans* [53,54]. This could also be the result of comparing the *Cx. restuans* tested in this study to a susceptible strain of a different species. The majority of *Cx. restuans* populations tested showed high susceptibility to malathion, so we wouldn’t anticipate increased levels in the biochemical assays. But there was one *Cx. restuans* population sampled from the Peoria region that displayed a high level of malathion resistance in the bottle assay (33% average mortality at the diagnostic time), but mosquitoes from this population did not display elevated levels of any detoxification enzymes or indication of insensitive ACHE. This could indicate the use of a different insecticide resistance mechanism, such as the thickening of the cuticle to reduce permeability to insecticides or increased metabolism of insecticides by elevated production of glutathione-S-transferases [55]. Assessing these other mechanisms was beyond the scope of the current study but would be valuable to explore in the future.

The *Culex* populations sampled in this study displayed great variation in their susceptibility to two insecticides commonly used in vector control, permethrin and malathion. The *kdr* point mutation (L1014F) was detected in half of the sequenced *Cx. pipiens* but was not found in *Cx. restuans*. Resistance in *Cx. pipiens* populations of IL. appears to be driven by a combination of *kdr* and enzymatic detoxification, while resistance in *Cx. restuans* is attributed to metabolic resistance alone. Further investigation is necessary to better understand the resistance status of *Culex* mosquito populations in IL. including the testing of additional populations, testing the susceptibility of other insecticides, and determining if there are additional mechanisms of resistance at play. This information is key to maintaining effective mosquito management strategies. Overall, this study highlights the necessity to implement mosquito insecticide resistance monitoring and management practices in Illinois.

## Acknowledgements

We are grateful for the technical assistance provided by Millon Blackshear, Corrado Cara, Erica Cimo, Katie Dana, Loyal Hall, Carly Kallembach, Andrew J. Mackay, Joseph Spina, Holly Tuten, and Jiayue Yan. We thank Janet McAllister for providing egg stock for the susceptible *Culex pipiens* colony “Chicago” strain. We also thank Aaron Baster from the Madison County Health District, Jeff Blackford from the Champaign Health District, Alan Bolds from the Wheaton Mosquito Abatement District, Brett Brosman from the Fayette County Health Department, Angie Crawford from the McLean County Health Department, Betty Croissant from the St. Clair County Health Department, Kristin Johnson & Sharon Verzal from the Kane County Health Department, Greg Maurice from the Dekalb County Health Department, Jason Probus from the Macon Mosquito Abatement District, Shelby Smith from the Adams County Health Department, Jack Swanson from the Peoria Health Department, Amber Wille from the Clay County Public Health Center, and Clark Wood from Clarke Mosquito, all for their time and willingness to assist with egg raft collections.

